# Global vascular plants reveal persistent gaps across taxa and ecoregions

**DOI:** 10.64898/2026.08.24.746674

**Authors:** Everton A. Maciel

## Abstract

Biodiversity aggregators such as GBIF provide unprecedented access to global biodiversity data, yet their representativeness remains uneven across space and taxa. This study examined the spatial and taxonomic structure of global vascular plant data available on GBIF. Six filters were applied to the GBIF vascular plant dataset, resulting in the removal of 54% of all records. Together, the filters explained more than 90% of the identified spatial issues, with duplicate and missing coordinates accounting for most of the variation. A higher number of occurrence records was associated with a greater number of spatial issues. Record distributions became progressively more even at finer taxonomic levels, from orders to species. The time series of occurrences for species, genera, and families increased sharply after 1800 and continued to rise, with no apparent stabilisation. Of the 824 ecoregions covered, 73 accounted for 72% of all occurrence records. These ecoregions spanned all continents but were strongly concentrated in Europe, followed by North America and Oceania. The analyses reveal four key patterns: (1) data volume is positively associated with spatial issues; (2) a small number of taxa account for a large proportion of records, whereas many are represented by relatively few; (3) occurrence data aggregated by GBIF have increased continuously since 1800; and (4) record coverage remains highly uneven across the world’s ecoregions. These results highlight the substantial contribution of biodiversity data aggregators to expanding access to biological information while demonstrating the persistent spatial and taxonomic biases that shape their contents. Such biases should be explicitly considered when assessing data completeness and quality and when using aggregated occurrence records to infer global biodiversity patterns.

**Highlights:**

- Spatial filtering removed 54% of global vascular plant records
- Retained data remained unevenly distributed across taxonomic levels
- Occurrence records increased sharply after 1800, with no apparent stabilisation
- Geographic gaps persisted across global ecoregions

## 1. Introduction

Open-access data platforms, such as the Global Biodiversity Information Facility (GBIF), which aggregate data from multiple biological collections, have become valuable sources of biodiversity data (Flemons et al., 2007; Petersen et al. 2021). GBIF’s mission is to make biodiversity information on species universally open and freely available via the internet (Lane and Edwards 2016). As of May 2025, GBIF was hosting over 113,000 datasets comprising more than three billion species occurrence records, documented by over 2,400 institutions (Sica et al. 2026). Many of these records originate from remote regions with high biodiversity, some of which are curated and stored in research institutions located in countries or even continents different from those where the specimens were collected (Edwards et al. 2000). This platform brings together biodiversity records from different regions and historical periods, extending the reach of collections beyond their physical holdings and enabling analyses across unprecedented spatial and temporal scales (Telenius, 2011; Zattara and Aizen, 2021). These records have been used in a wide range of scientific fields, with research based on GBIF data in macroecology leading the way over the past 17 years, alongside studies on taxonomic decline, species interactions, and disease ecology (Heberling et al. 2021). Given that taxonomy is a planetary science requiring a planetary-scale tool, GBIF is well placed to fulfil this role (Wheeler et al. 2004). In this context, GBIF guides ongoing initiatives and the strategic prioritisation of biodiversity data aggregation across diverse fields of knowledge, including environmental sciences and policy, evolutionary biology, conservation, and human health (Heberling et al. 2021).

The potential use of aggregator platforms such as GBIF depends on the quality and representativeness of the available data, as the inherent limitations of these datasets encompass both information gaps and issues related to the accuracy and representativeness of records (Anderson et al. 2020). Therefore, accessing aggregated datasets from GBIF requires know-how to access specimen records properly (Folk and Siniscalchi 2021; Steinke et al. 2025) and to account for the gaps and biases associated with them. Regarding bias, not all existing data are suitable for use, as many records lack geographic coordinates, are assigned to the ocean (Colli Silva et al. 2020; Führding Potschkat et al. 2022), or are erroneously assigned to particular locations (Maldonado et al. 2015). Strong taxonomic and geographic biases are also created by data voids, with well-studied groups such as birds being over-represented while insects, fungi, and non-tree plants are neglected, and records being concentrated in accessible areas (Feeley 2015; Farooq et al. 2021). For instance, these records exhibit persistent spatial biases towards roads, herbaria, urban centres, accessible areas, and lower elevations (Botts et al. 2011; Daru et al. 2017; Oliveira et al. 2025). Although conservation strategies depend on species records in many aspects, species are going extinct faster than these issues can be solved (Pimm et al. 2006).

Regarding gaps, the “data void,” which refers to the scarcity of digitised and accessible species occurrence records, is one of the major limitations associated with aggregation platforms such as GBIF (Feeley and Silman, 2011). This gap contributes to the Wallace shortfall by limiting our knowledge of species distributions, especially in higher-diversity regions (Lima et al., 2024). This lack of data may limit the assessment of rare species and their protection status (Maciel and Martins, 2021), as well as the prediction of species’ niches in response to future climate change, since both strategies depend on a reasonable understanding of species distributions (Beck et al., 2014; Qiao et al., 2017). Data gaps affect a vast proportion of the globe, particularly species-rich regions, rendering part of biodiversity functionally invisible (Trindade and Marques, 2024; Feeley, 2015). For instance, 74% of South American plant species have fewer than 20 records, and 10% have no digitised records whatsoever (Colli-Silva et al., 2020; Trindade and Marques, 2024; Feeley, 2015). Consequently, global datasets contain substantial gaps, leaving some regions of the globe with blank spaces, particularly in areas of high biodiversity (Ronquillo et al., 2023; Blades et al., 2025).

Due to biases and data voids associated with data from aggregator platforms, actions are required to improve data quality and make records more suitable for specific purposes (Veiga et al., 2017; García-Roselló et al., 2015). This requires, on the one hand, quality control to prevent and correct errors and, on the other, ensuring that the selected data meet the quality standards required for a particular purpose (Veiga et al., 2017). This process involves filtering and excluding data that do not meet the necessary quality criteria, with missing coordinates (Maciel et al., 2021) and duplicated records (Dorey et al., 2023) being among the most common problems detected. As manual data cleaning is often impractical for large volumes of data, automated routines offer an alternative approach to overcoming these limitations (Führding-Potschkat et al., 2022). Automated workflows provide various filters to address issues such as duplicate records, collection year, and coordinates located within urban areas or in the sea for terrestrial taxa, among others (Zizka et al., 2020). In the case of data gaps, one alternative is to combine data from multiple sources in an attempt to improve the information available in regions where data are extremely scarce (Heberling et al., 2021; De Araujo et al., 2022; Maciel and Guilherme, 2023). However, in practice, the prevailing approach to data gaps has been to identify where they occur and treat them as limitations (Maciel and Martins, 2021; Xie et al., 2025). Thus, improving the quality of occurrence records from biological collections remains a challenge, particularly for diverse groups such as vascular plants (Chapman, 2005).

Vascular plants (Tracheophyta) are one of the key groups of organisms in terrestrial ecosystems, playing a fundamental role in habitat structure, primary productivity, and nutrient cycling (Begon and Townsend, 2020). The GBIF global vascular plant dataset contains 440,562 accepted names, totalling 1,441,152 occurrence records (Govaerts, 2026), which have been used to address a wide range of questions across different spatial scales and fields (Heberling et al., 2021). In this context, global plant databases are at the centre of ongoing discussion. This discussion is motivated, on the one hand, by the transformation that herbaria are undergoing in the age of big data, driven by the ambition to make available information on the plant world accessible online (Folk and Siniscalchi, 2021; Davis, 2024). On the other hand, the debate is fuelled by the use of data already available, with biases associated with these records and spatial gaps resulting from data gaps being key issues (Rocha-Ortega et al., 2021; Beck et al., 2014; Troudet et al., 2017).

This study addresses this knowledge gap by systematically evaluating the impact of six distinct spatial filters applied to the global vascular plant occurrence dataset mobilised through the Global Biodiversity Information Facility (GBIF). The effects of the filters were examined at the dataset level, across taxonomic levels, and across global ecoregions. At the dataset level, each filter’s contribution to coordinate removal was quantified as a proportion of the total number of GBIF records. The remaining records were analysed across orders, families, genera, and species to assess taxonomic coverage. Their temporal patterns were quantified by tracking occurrence counts at each taxonomic level through time. Finally, their spatial patterns were examined based on their distribution among global ecoregions. By quantifying how different filters affect biological data quality, this study provides a methodological basis for identifying global geographic and taxonomic gaps (Pitogo et al., 2026; Wilcox et al., 2025). This diagnostic tool can guide future surveys towards historically data-poor regions and taxa (Hughes et al., 2021), providing the empirical basis needed to support global biodiversity governance and conservation (Sterner et al., 2020).

## 2. Method

### 2.1. Data sources and downloaded

The classification of Tracheophyta, as well as the taxonomic backbone used by GBIF (Bánki et al., 2026), was adopted for this study. Tracheophyta records were retrieved via the Application Programming Interface (API) (Wieczorek et al., 2026). APIs are methods that promote the exchange of data between independent systems, enabling researchers to access billions of occurrence records without manually downloading files from the web portal (Norton, 2021). The rgbif package was used to facilitate the download of GBIF data through the API (Chamberlain et al., 2017). To apply download functions through rgbif, the researcher must provide their credentials (username, password, and email; see Chamberlain et al., 2022), while the GBIF API handles the creation of the DOI and the asynchronous processing of the file (Vasconcelos and Boyko, 2025). Once the credentials had been set up and the authenticated download request had been submitted, the next step was to apply the name_backbone() function to extract the taxonomic key (taxonKey) for Tracheophyta from GBIF. This taxonomic key was then used as a search criterion in the occ_download() function to allow all occurrence records linked to the Tracheophyta taxonKey in GBIF to be downloaded via the GBIF API. The processing of the request was monitored using the occ_download_wait() function. Once complete, the occ_download_get() function was used to retrieve the file containing the records, while the occ_download_meta() function was used to access the request information and metadata (Supplementary 1).

**Figure 1.**
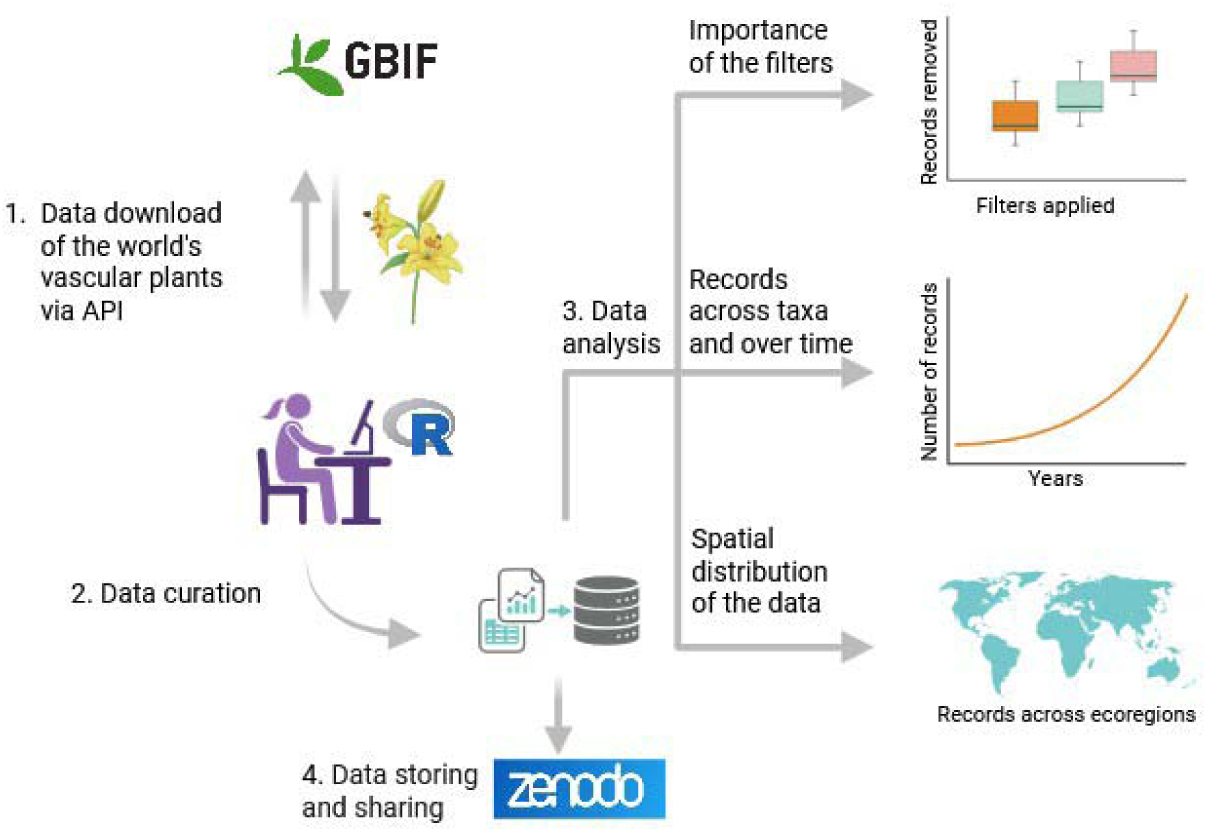
Study workflow diagram.

Since the Tracheophyta dataset combined records from multiple families into a single large file, the dataset was split into family-level files. This facilitated data processing and subsequent analysis, even on a low-performance computer. The archive and data.table packages were used to manipulate these files, as they provide efficient tools for handling large datasets (Dowle et al., 2019). First, the archive_extract() function was used to extract the files from the Tracheophyta.zip file. Next, the fread() function from the data.table package was used to import the occurrence file, selecting only the relevant columns and specifying the data type of each variable in advance. This approach reduced memory usage and improved computational performance. After importing, records lacking geographical coordinates or family identification were removed. The result was a temporary data frame in R’s memory for each family, containing only the selected columns (see Supplementary 2, Tables 1 and 2). The raw Tracheophyta data retrieved from GBIF can be accessed through the GBIF DOI (see GBIF, 2026). These data were subjected to a filtering process to remove spatial artefacts (see next section).

### 2. 3. Spatial and Geographic curation

The records were submitted to a spatial cleaning process using the CoordinateCleaner, an r-package to scan datasets of species occurrence records for geo-referencing, tailored to problems common in biological and palaeontological databases and can handle datasets with millions of records (Zizka et al. 2019). Although this package have numerous filters to detect flags, only six of them were applied here, namely cc_dupl() for remove duplicate records, cc_equ() for identical coordinates, cc_gbif() for points near the GBIF headquarters in Copenhagen, cc_inst() for records associated with biological institutions and cc_sea() for records located in marine environments incompatible with the study group. The filters were applied for each family separately, with the number of records removed by each filter being recorded for subsequent analysis. Processing was carried using the data.table package to efficiently import and export data via the fread() and fwrite() functions, respectively (Dowle et al. 2019) (Supplementary 3). The final datasets for each family were organized into single folders, compressed into ZIP files, and uploaded to Zenodo in Comma-Separated Values format (.csv). The datasets for each family were also converted to geospatial data GeoPackage (.gpkg) format using the R package sf (Pebesma 2018), organized into single folders and compressed into ZIP files. The cleaned datasets in both formats can be accessed in Zenodo (Maciel 2026).

### 2. 3. Data analyses

#### 2. 3. 1. Data flags

To assess the effect of spatial issues on all dataset, the proportion of records removed was calculated by each filter relative to the total number of records available for each family in GBIF. The proportions of records for each filter were then used as the response variable (Y), and the filters as categories for built boxplots were to compare the extent of record removed among filters. This was done in order to analyze the magnitude of the effect of each filter on the initial volume of data records, the number affected by spatial issues, and the amount of data retained after quality filtering. A Pearson correlation matrix was calculated using the absolute value of the six filters to assess the degree correlation among them. A Principal Component Analysis (PCA) was performed in R using the prcomp() function to analyse the main gradients of variation in the data flags recovered from cleaning process (Supplementary 5). The six filters were centred and standardised prior to analysis. The proportion of total variance explained by each PCA axis was calculated from the corresponding eigenvalues. For each principal component, the corresponding eigenvalues, proportion of explained variance and cumulative explained variance were obtained. The filters loadings for each component were then used to identify the variables most strongly associated with the main axes of ordination. Subsequently, regressions were fitted between the logarithm of the total number of records available download from GBIF (X) with both the number of records removed and retained after filtering (Y) for families (Supplementary 2, Table 3 and 4).

#### 2. 3. 2. Taxonomy dimension

Occurrences for each taxon were aggregated by summing the total number of records across the entire time series for each taxonomic level (order, family, genus, and species). The relative contribution of each taxon was then calculated as a proportion of the total number of records. Taxa were ranked in descending order of relative abundance, and cumulative percentages were calculated. At the genus and species levels, only taxa whose cumulative contribution reached the 50% threshold were retained for visualisation, while the remaining taxa were grouped as ‘others’ and excluded from the figure. This exclusion was necessary because there were too many species and genera to be interpretable in a single figure. A treemap was generated to represent the relative contribution of the dominant taxa, with the area of each tile proportional to the taxon’s percentage of the total number of records (Supplementary 4). The figure was produced using the ‘ggplot2’ and ‘treemapify’ packages in R (Wickham et al., 2009; Wilkins, 2021, respectively).

#### 2. 3. 2. Temporal and spatial dimension

For each taxon, a data matrix with two columns (taxon name and year) was constructed. The data for each taxon were then grouped by year using dplyr (Wickham et al., 2023), and the number of distinct taxa observed per year was calculated using the n_distinct function. The resulting trend was visualised using ggplot2 as a connected scatterplot with a locally estimated scatterplot smoothing (LOESS) curve (span = 0.75) and its 95% confidence interval. The LOESS curve was used to reduce the influence of short-term fluctuations among individual observations and highlight the overall trend in the data without assuming a linear relationship. The 95% confidence interval provides an indication of the uncertainty around the estimated trend. For the analysis of spatial trends, only the number of families was considered, as this metric provides a consistent measure of taxonomic representation while reducing the influence of differences in the number of species or records among families (Supplementary 6).

Each family file (.gpkg) was spatially intersected with the ecoregion layer, which classifies 867 ecoregions (Olson et al., 2021), to identify the ecoregion in which each occurrence was located. The number of records for each ecoregion was then calculated by counting the occurrence points falling within its boundaries. The proportion of records represented by each ecoregion was then calculated relative to the total number of occurrences retained in the final dataset. This analysis was not performed at all taxonomic levels, as the focus was on the overall distribution of records across ecoregions, providing an indication of the contribution of each ecoregion to the total volume of records. To visualise the global distribution of records, the absolute number of occurrences per ecoregion was categorised into 20 classes using the natural breaks method and displayed on a map (Supplementary 6).

To assess the relationship between records and latitude, a Generalized Additive Model (GAM) was employed, with the gam() function from the mgcv package in R fitted to the data. Due to the highly skewed of the data, the records were transformed using natural logarithm of 1 + x (log1p) to reduce skewness and minimize the influence of unusually high values. Latitude was expressed as the absolute distance from the Equator (abs(Lat)), enabling us to evaluate the latitudinal gradient independently of hemisphere. This variable was included in the model as a smooth term to allow for a potentially non-linear relationship between species richness and distance from the Equator. Sampled area was included as a covariate and was also transformed using log1p(area_km²) to account for the effect of spatial extent among sampling units. The model was fitted using restricted maximum likelihood (REML). The significance of the smooth term was assessed using the associated non-parametric test. The shape of the latitudinal relationship was visualised using model predictions across a gradient of distance from the Equator while holding the ecoregion area constant at its median value. The peak in data records was defined as the distance from the Equator corresponding to the maximum records along the fitted curve (Supplementary 7).

## 3. Result

### 3.1. Data flags

Around 54% of the original records downloaded from GBIF were discarded. Most of the removed data were due to duplicate or missing coordinates (Figure 2A). The Pearson correlation revealed strong correlations among several filters, as shown by duplicates and missing coordinates (r = 0.95), equal latitude and longitude and missing coordinates (r = 0.94), duplicate and biodiversity institutions (r = 0.89), and missing coordinates and biodiversity institutions (r = 0.89), while records around Copenhagen showed weaker positive correlations with all filters (Figure 2B). PCA revealed a strong multivariate gradient, primarily represented by the first principal component (PC1), which explained 79.11% of the total variance (Figure 2C). Together, PC1 and PC2 explained 90.26% of the total variance, capturing most of the variation in the data filters. PC1 was mainly associated with missing coordinates, duplicates, biodiversity institutions, equal latitude and longitude, and sea and coastal lines, indicating that these filters varied together and were the main factors contributing to this component. PC2 was primarily related to records around Copenhagen, with a secondary contribution from Sea and coastal lines. The Pearson correlation analysis supported these patterns, showing strong relationships among several filters. Finally, there was a positive correlation between the absolute number of records removed and the number of records retained after filtering, as well as the total number of records available in GBIF, indicating that greater data volume was associated with an increasing number of spatial issues (Figure 2D).

**Figure 2.**
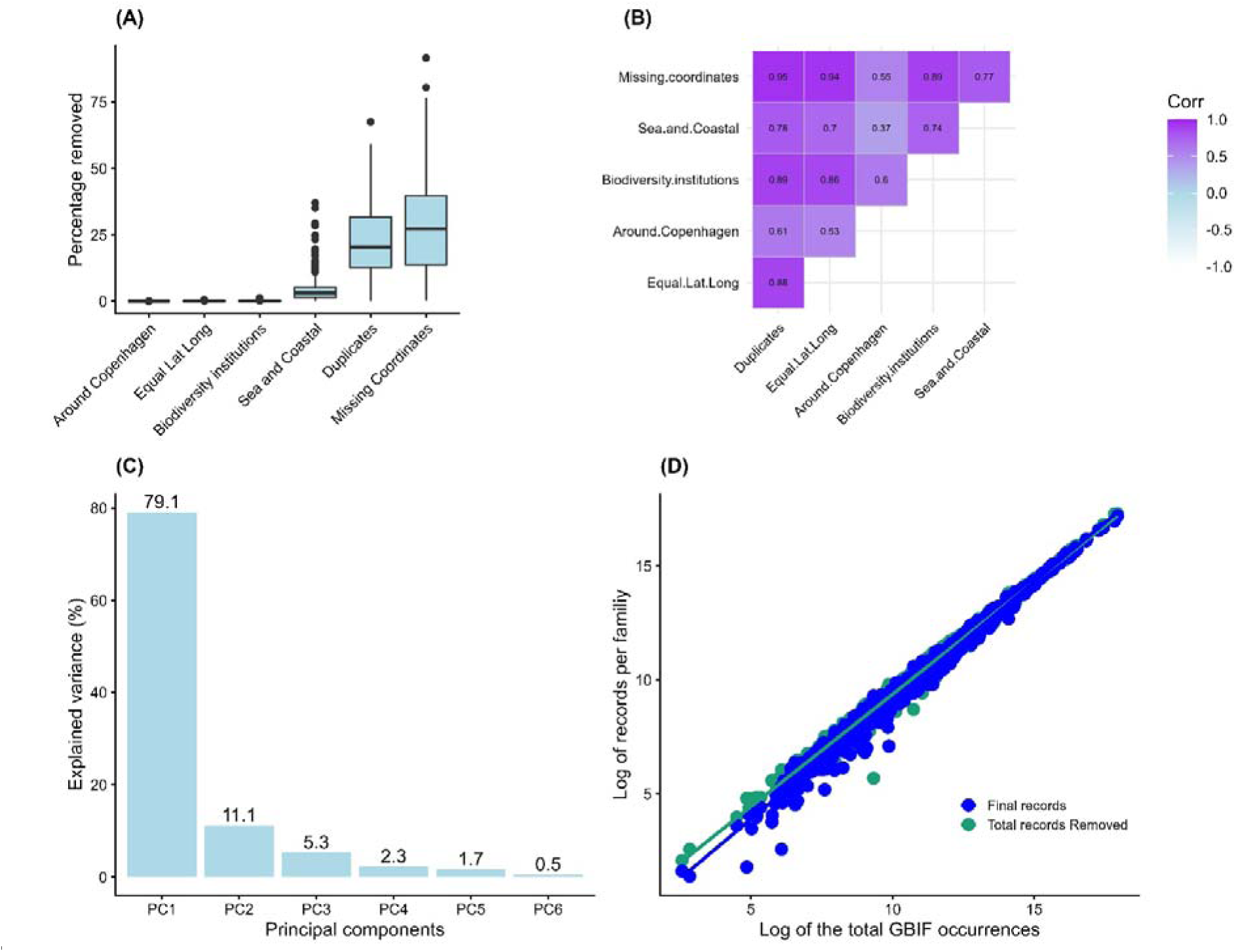
Data flags and patterns of removed data. Percentage of GBIF occurrences among filters (A). Correlation matrix among the six filters used to clean GBIF occurrences (B). Principal components (PCs) showing the five components, with the percentage of explained variance indicated at the top of the bars (C). Scatter plot illustrating the correlation between the log-transformed total number of GBIF occurrences and the number removed by filters, with the log-transformed number of occurrences retained in the final database (D). Each point in the scatter plots represents a family (n = 461).

### 3. 1. Taxonomy dimension

The final dataset comprises 28,285,682,100 records, representing 46.5% of all gross data downloaded from GBIF. It contains data on eight classes, 85 orders, 461 families, 15,357 genera and 307,674 species. The number of cleaned occurrences among taxa ranges from 29,503 (Cycadopsida) to 214,515,319 (Magnoliopsida) at class level, from 100 (Petrosaviales) to 38,299,910 (Poales) at order level, from 4 (Petiveriaceae) to 29,278,105 (Asteraceae) at family level, from 1 (n = 814) to 6,652,353 (*Carex*) at genus level, and from 1 (n = 36,577) to 939,710 (*Urtica dioica*) at species level. The proportion of occurrences that each taxon represents relative to the total number of records in the dataset also varies considerably, with only a few taxa accounting for a relatively large proportion of the records, while most were represented by very low proportions (Figure 3). The highest taxonomic levels showed the strongest concentration of records, with one taxon accounting for around 13.5% of all records at order levels, while the most abundant family represented around 10.4% of total occurrences. This pattern became less pronounced at lower taxonomic levels, with 2.4% and 0.3% at genus and species level, respectively.

**Figure 3.**
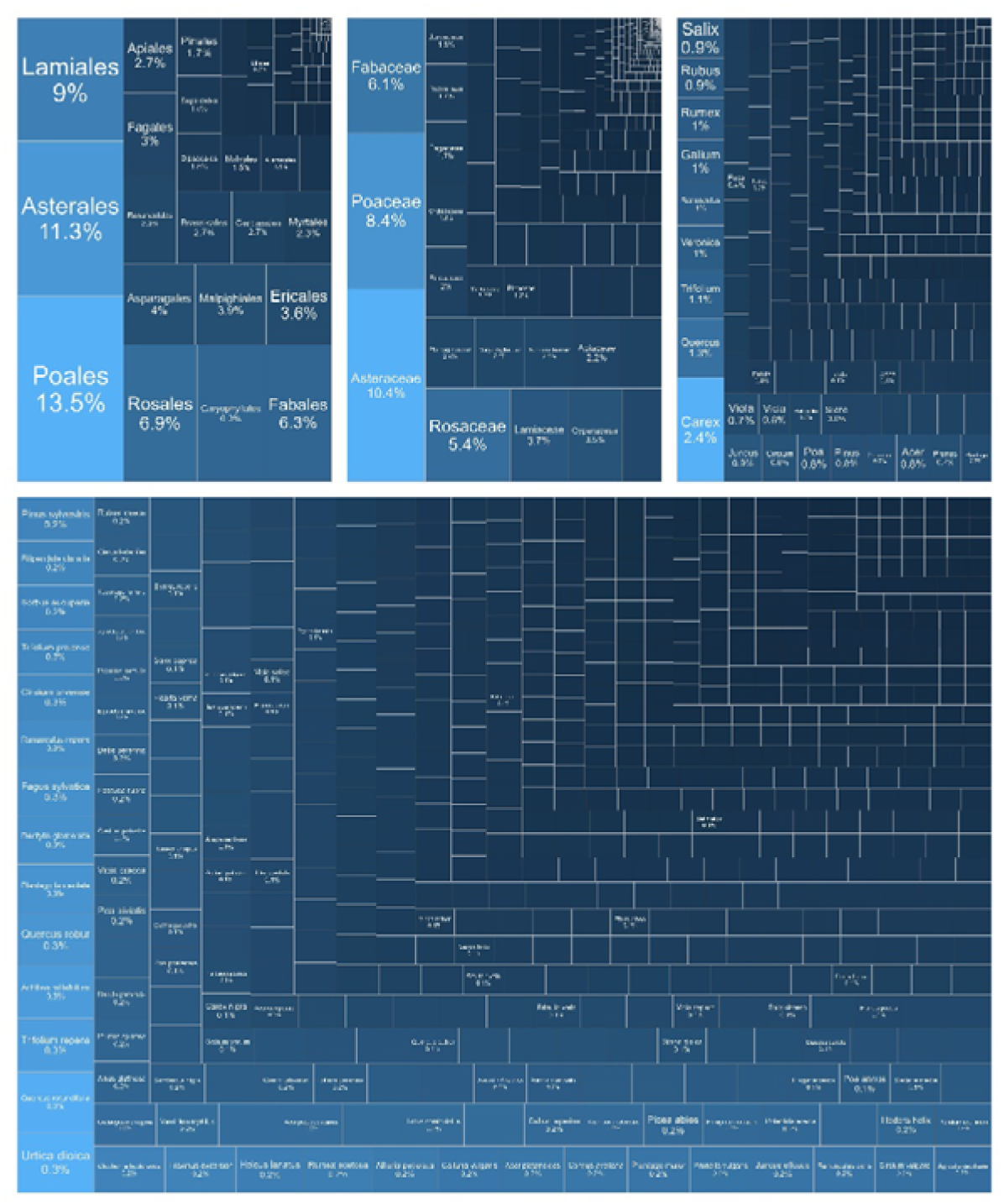
Treemap showing the relative contribution of the dominant taxa to the total number of records. Only species accounting for the top 50% of cumulative occurrences are displayed. Tile area is proportional to the percentage of total records represented by each taxon.

### 3. 2. Temporal and spatial dimension

The time series of order level records displays a sigmoidal accumulation trajectory, with a modest initial development phase, a notable expansion beginning in the nineteenth century, and a gradual decline in the growth rate in more recent decades (Figure 4A). The time series of family, genus, and species level records, on the other hand, shows a steady increase throughout the study period, with a noticeable acceleration after the middle of the nineteenth century and no obvious signs of saturation by the end of the record (Figure 4B–D).

**Figure 4.**
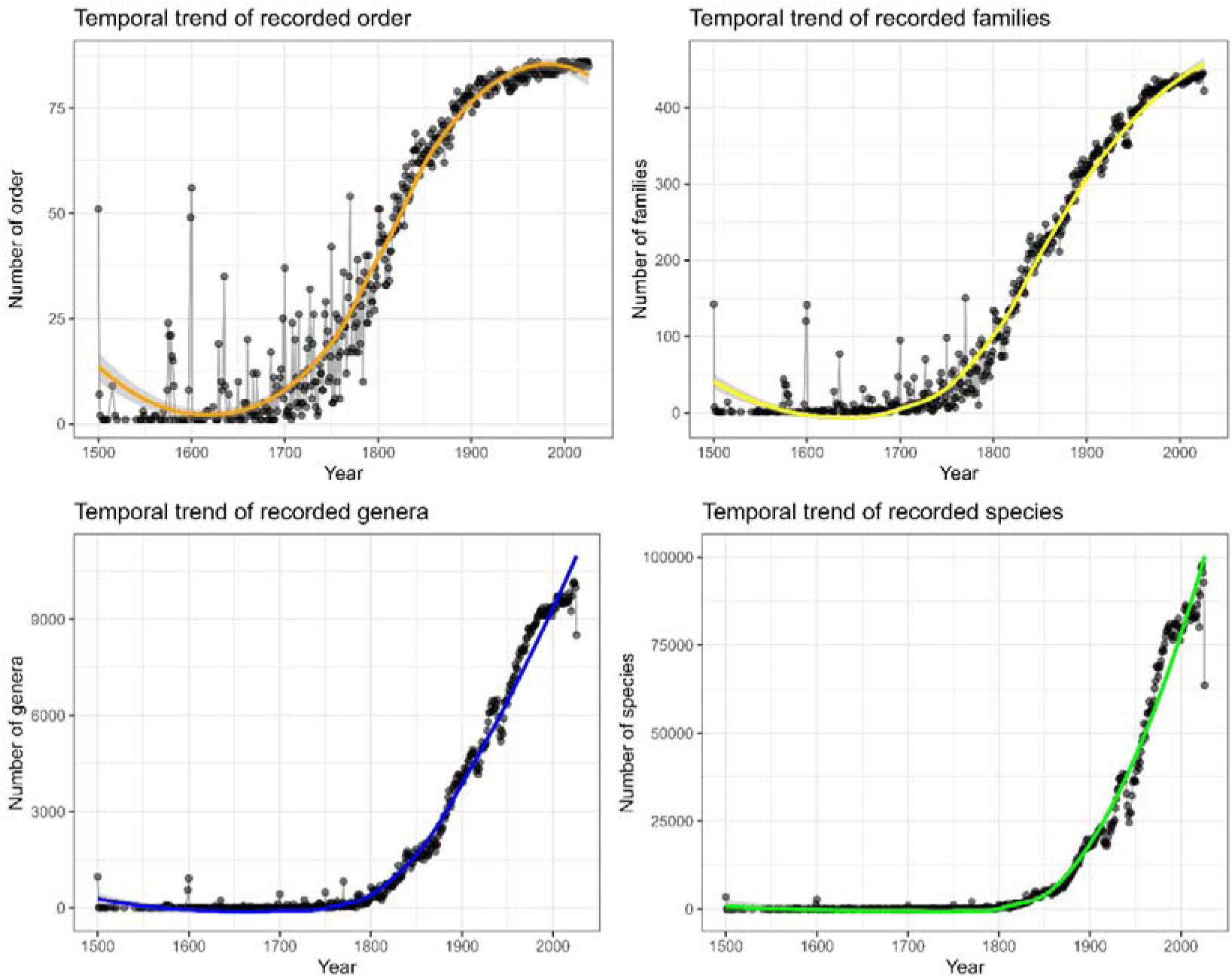
Temporal trend of GBIF-recorded taxa over time. Annual data points are connected by lines to illustrate year-to-year fluctuations, while the LOESS smoothing curve (span = 0.75) with its 95% confidence interval (green line) shows the long-term trend in the number of taxa observed over time. A = order, B = families, C = genera, and D = species.

The data occurrence covers 827 ecoregions, in which the number of records varies greatly across them (Figure 4; Supplementary 2, Table 4). Two ecoregion groups can be distinguished in terms of total records inside each of them. The first group comprising 73 ecoregions holds more than 0.15% of all records each and collectively accounts for 82.44% of all records (Figure 4A). These ecoregions were distributed amongst the regions as follows: Europe (41), North America (27), Oceania (9), Asia (4), Africa (3) and South America (3), respectively. The second is a larger group comprising 754 ecoregions (with 0.15% or less of all records) together accounts for 17.56% of all occurrences (Figure 4A). In both groups, records are unevenly distributed among ecoregions. For example, in the first group 20% of all records are concentrated in two ecoregions, Atlantic mixed forests and Western European broadleaf forests, 13 ecoregions contain between 1.15% and 7.6% of all records, and 25 contain between 0.16% and 1%. The second group includes only five ecoregions with 0.15% of the records, while the massive remainder number of ecoregions (749) contains less than 0.15% each.

GBIF records exhibited an asymmetric distribution in the original data, with a higher concentration of low values and a tail of high values (Figure 5). Following logarithmic transformation, the distribution of records became approximately normal, reducing the asymmetry and providing a more suitable distribution for spatial modelling (Figure 5C). The log-transformed number of records across ecoregions showed a significant and nonlinear relationship with distance from the equator (EDF = 2.32; P = 0.0015; Supplementary 2, Table 5), indicating that the relationship between records and latitude cannot be adequately described by a simple linear effect. The GAM fitted curve showed a unimodal pattern, with the number of records increasing towards intermediate latitudes and reaching an estimated maximum at approximately 49.3° from the Equator (Figure 5B). Ecoregion area (km²) also had a positive and significant effect on the number of records, indicating that larger ecoregions contained more records (β = 0.0333 ± 0.0054; z = 6.21; p < 0.001).

**Figure 5.**
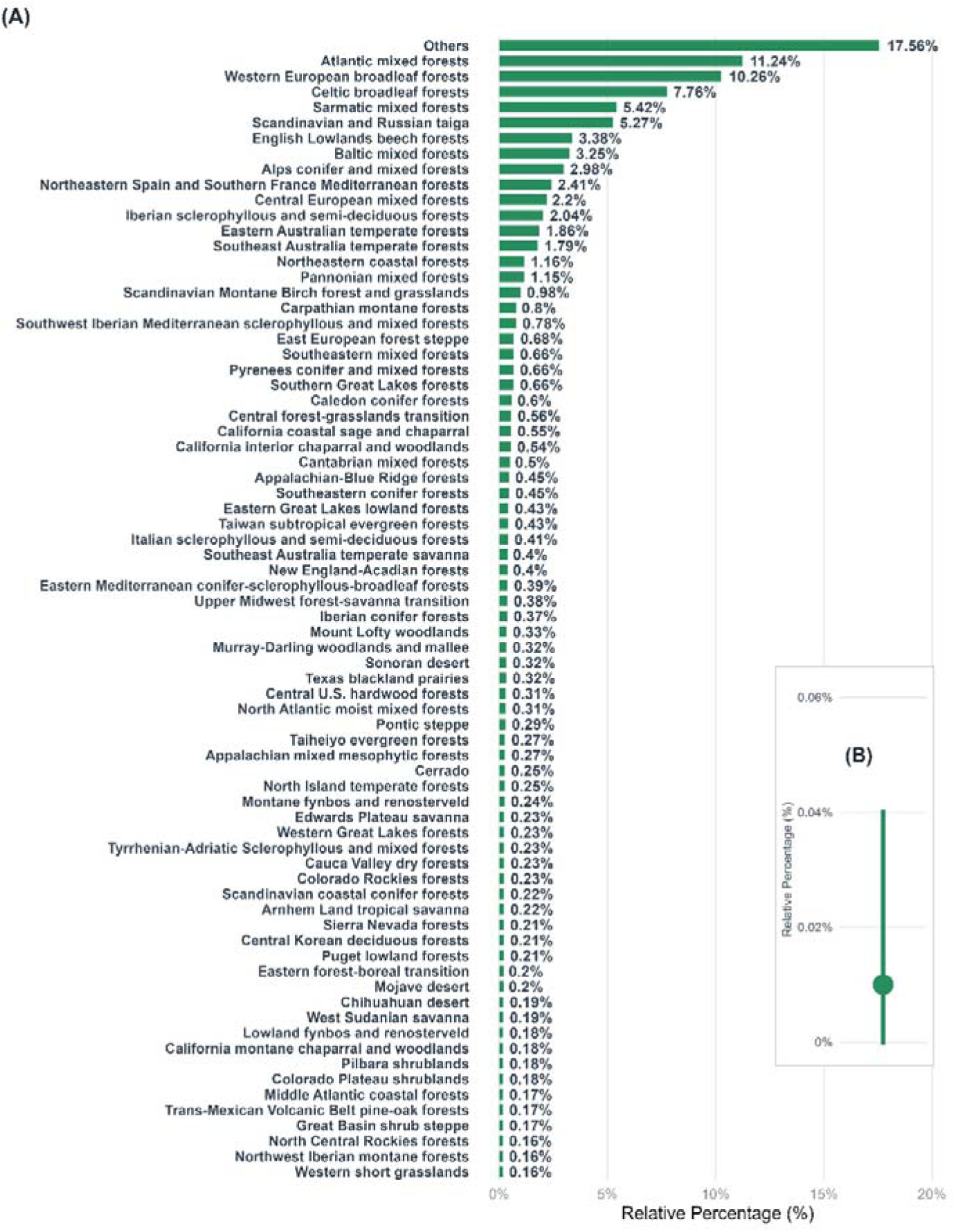
Percentage of GBIF occurrences by ecoregion. The bars and numbers represent the percentage of occurrences relative to the total number of occurrences for the 75 ecoregions with the highest values, while ‘others’ represents all other ecoregions with less than 0.15% of occurrences (A). The median and standard deviations are shown in relation to the proportion of occurrences of each ecoregion across the entire dataset (B).

**Figure 6.**
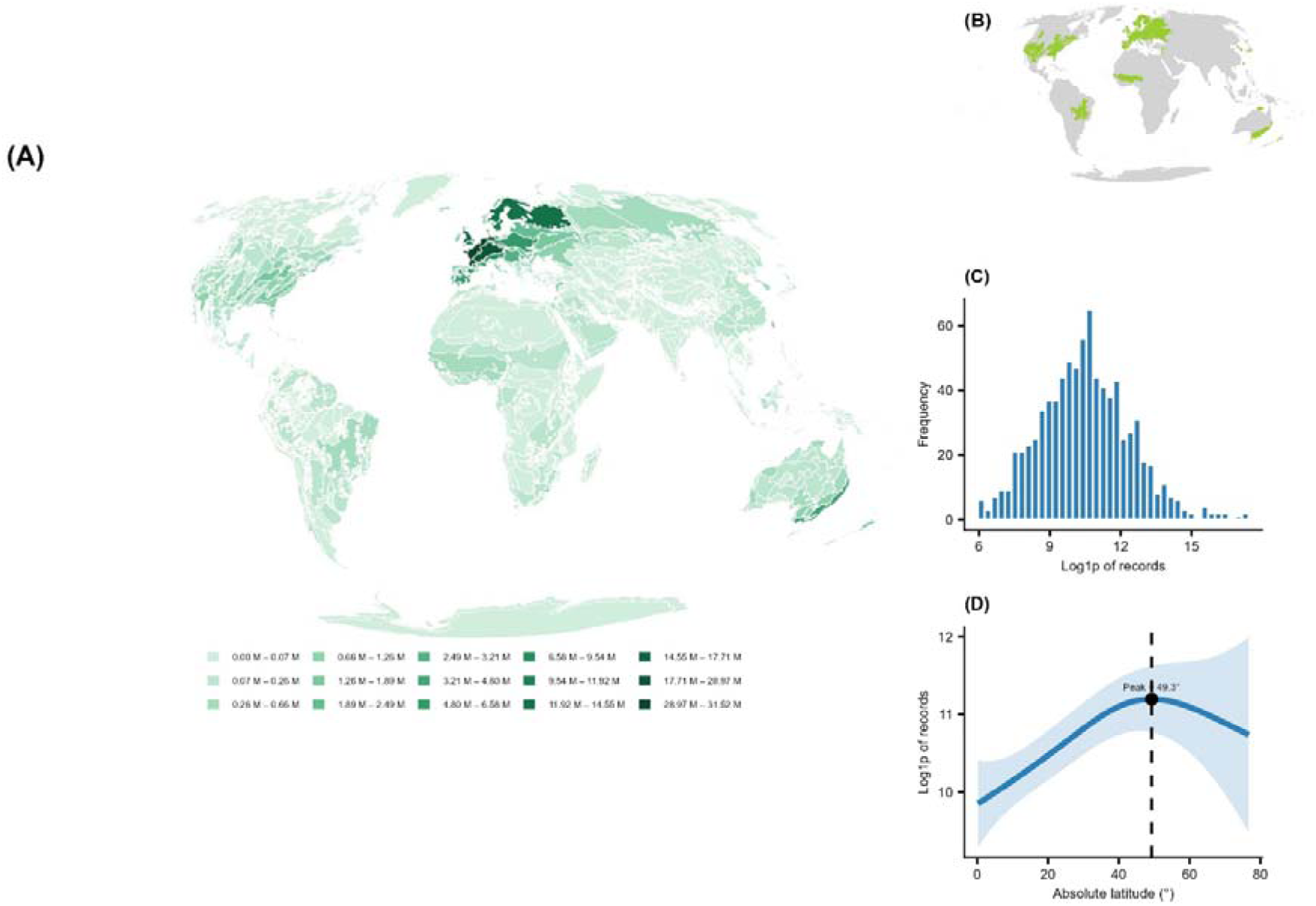
Distribution of occurrences across ecoregions (A). The map legend represents 20 natural-break classes applied to the absolute number of occurrences in the ecoregions. The smaller map shows 73 ecoregions, which account for 82.44% of all occurrences (B). The bar chart shows the distribution of occurrences following the log1p transformation (C). The graph shows the GAM fit to the log1p-transformed number of occurrences and log1p-transformed absolute latitudes, with the white point indicating the peak number of occurrences (D).

## 4. Discussion

The Global Biodiversity Information Facility (GBIF) has become a major resource for biodiversity science in the era of big data. By aggregating biodiversity data from a variety of datasets across the world, GBIF has expanded access to information on species occurrences, enabling biodiversity assessments at local, regional, and global scales, as well as supporting research, monitoring, and conservation. The unprecedented volume and variety of these data also provide an opportunity to examine not only where biodiversity records are located, but also how consistently they represent the distribution of the taxa, regions, and time periods from which they originate. The present results reveal that, despite the substantial quantity of available records, their usability and representation vary considerably across biogeographic dimensions. This variation results from substantial data loss during quality filtering, as well as spatial and temporal variability. These findings do not diminish the value of GBIF but rather highlight the need to understand patterns of data availability and quality across space and time to maximise the scientific value of large biodiversity datasets for research and conservation.

### 4.1. Data flags

Although the use of filters is necessary to ensure spatial consistency and reduce potential biases associated with missing or duplicated coordinates, the removal of approximately 54% of the original records inevitably reduced the amount of information available for subsequent analyses. The data loss resulting from spatial filters is strongly precedented; for instance, Eckert et al. (2024) discarded 38.8% of vascular plant records (4.7M out of 12.2M) when filtering georeferences with high coordinate uncertainty. This trade-off highlights a deeper epistemological challenge: the "datafication" and standardization of biological records to fit digital formats often strip away local biophysical context, reducing the multi-dimensional ecological niche to a simplified, non-interacting "data niche" (Devictor and Bensaude-Vicent 2016). Furthermore, centralized aggregation infrastructures such as GBIF often fail to accommodate localized taxonomic variations, applying globally homogenized classifications that can exclude valid biological information (Eckert et al. 2024). These taxonomic and spatial omissions have severe macroecological consequences, as severe data gaps can distort downstream analyses to the point of completely inverting modeled species-climate relationships (Quian et al. 2022). Therefore, overcoming global biodiversity shortfalls requires filtering approaches that carefully balance quality control with geographic and taxonomic representation, ensuring that conservation priorities are guided by a reliable evidence base (Hughes et al. 2024; Pitogo et al. 2026; Wetzel et al. 2028).

While GBIF provides a large volume of records that may initially suggest broad sampling coverage, a substantial proportion of these may be unsuitable for spatial analyses after quality control. Although this substantial reduction represents a clear trade-off between data quantity and reliability (Ronquillo et al. 2023), retaining records with spatial inconsistencies could introduce greater uncertainty into subsequent analyses (Maldonado et al. 2015). Conversely, inaccurate or poorly georeferenced records can obscure the actual distribution of populations, potentially affecting estimates of range size and, ultimately, assessments of conservation status (Panter et al. 2020). Cleaning the data therefore serves not simply to reduce noise, but to constrain spatial analyses to records with a defensible geographic basis. The importance of these filters is further supported by the multivariate structure of the quality flags. The first principal component accounted for 79.11% of the total variance and was strongly associated with missing coordinates, duplicate records, biodiversity institutions, equal latitude and longitude, and records associated with sea or coastal lines. This indicates that these sources of uncertainty were not independent problems, but rather formed a broader gradient of spatial data quality. Although many other filters are available, the filters related to the total number of records removed accounted for most of the variation explained by the first two PCA axes, which together explained 90% of the total variance. This suggests that they are important indicators of spatial issues in GBIF data.

### 4.2. Taxonomy dimension

Taxonomic curation can substantially alter the structure of biodiversity records, both by addressing spatial issues and by changing the distribution of records across taxonomic levels. The occurrence records after issued removed varies greatly among order, families, genera and species. For example, in family the proportion of records removed varied from 2.5 to 97%, indicating notable heterogeneity in data quality and spatial completeness across taxonomic groups. In this study, the most represented family accounted for 10.4% of all cleaned records, compared with 2.4% for the most represented genus and only 0.3% for the most represented species, indicating a marked decline in record concentration as taxonomic resolution increases. This pattern is consistent with the broader effects of taxonomic cleaning, which can substantially reduce the number of accepted names while retaining a highly uneven subset of occurrence records; as observed in the flora de Bogotá, which this process reduced the number of family and species names by 24% and genera by 7% (Vargas et al. 2024). Such filtering can therefore amplify pre-existing taxonomic imbalances in occurrence databases, including GBIF, where most records are identified to species but identification accuracy varies considerably among taxa (Svenningsen and Schigel 2024). Consequently, differences in record availability should not be interpreted uncritically as signals of taxonomic importance, abundance, or diversity, but as patterns shaped by both biodiversity and the uneven processes through which taxa are collected, identified, curated, and represented in the database.

The positive correlations between the total number of GBIF records and the number of records retained and removed show that greater data availability does not necessarily indicate higher data quality. Large datasets provide more records for analysis, but they also contain a greater absolute number of records affected by spatial, taxonomic, or other inconsistencies. This pattern is supported by previous studies, in which data-cleaning procedures removed substantial proportions of GBIF records, including 87–91% of a large Brazilian plant dataset (Ribeiro et al. 2022), 31.7– 62.7% of records in a well-documented genera of plant (Führding-Potschkat et al. 2022) and about 50% of beetle records (Blades et al. 2025). Based on an analysis of the reduction in GBIF data volumes for well-documented taxa, these studies collectively support the evidence found in these studies that large data volumes come at the cost of increased spatial noise. The number of GBIF records should therefore not be treated as a direct proxy for the amount of reliable spatial information available, reinforcing the need for rigorous filtering and quality control before spatial analyses.

### 4. 3. Spatial and temporal dimension

The concentration of records in ecoregions primarily located in Europe, North America and southern Australia further suggests an uneven geographical distribution of biodiversity data availability. These results support previous findings, which found that 79% of all GBIF occurrence data comes from just 10 countries, predominantly concentrated in Europe, the USA and Australia, leaving only 18% for the other 240 countries and territories (Hughes et al. 2021). While some ecoregions may contain substantial biodiversity, their high representation in the dataset is unlikely to be solely due to differences in biodiversity and may also be influenced by variation in sampling effort, accessibility, research activity and the availability of digitised occurrence data. In fact, the geographic and taxonomic coverage of digitally accessible data is disproportionately high in North America, Western Europe, and parts of Australia (Meyer et al. 2015). Conversely, the relatively low representation of several ecoregions in the Global South may indicate important spatial data gaps, particularly in regions that could be biologically significant for the considered taxa. In some regions, species richness was strongly correlated with the number of records, leading to the conclusion that species richness might be an artefact of data recording (Yang et al. 2013). Many regions have been extensively documented, whereas others remain comparatively poorly understood. The extent to which these uneven patterns of data availability distort our understanding of biodiversity remains unclear, particularly because many studies rely on data from open-access platforms that are themselves affected by substantial spatial and temporal sampling biases.

A small number of ecoregions accounted for a disproportionate share of records, whereas many others were represented by very few observations, with some contributing ≤0.15% of the total. This unevenness reflects not only differences in where biodiversity occurs, but also where, how intensively, and under what conditions it has been documented (Meyer et al. 2015; Feeley et al. 2015). In particular, well-sampled regions and grid cells are disproportionately concentrated in high-income countries, consistent with the influence of economic capacity on research effort and the production, digitisation and accessibility of biodiversity data (Ronquillo et al. 2023, Blades et al. 2025, Hughes et al. 2021). The Afrotropical distribution of *Catharsius* records illustrates this disparity: records are concentrated largely in northeastern South Africa and Kenya, while extensive and highly diverse areas of Central and West Africa remain poorly represented (Blades et al. 2025). Thus, the apparent concentration of biodiversity records is partly a record of where research has been conducted and data have become available, rather than a direct representation of the spatial distribution of biodiversity.

Previous studies have shown that highly species-rich ecoregions in South America, including the Caatinga, Cerrado and Madeira-Tapajós moist forest, remain affected by substantial gaps in occurrence data (Feeley et al. 2015). This reflects a data paradox, which states that hotspots of biodiversity remain poorly represented in biodiversity databases, as recently demonstrated for the Atlantic Forest (Trindade and Marques, 2024, Colli-Silva et al. 2020). Against this background, the Cerrado emerged as interesting pattern, despite having comparatively lower plant diversity than the ecoregions along the Brazilian coast and in the Amazon, ranked among the 75 ecoregions with the highest proportion of occurrence records (> 0.15%). This relatively high record volume should not, however, be interpreted as evidence of adequate sampling. Given the vast extent of the Cerrado, its maybe remains very low, since previous estimates indicating approximately one collection per 30 km² (Feeley et al. 2015). Conversely, even intensively collected regions such as the Atlantic Forest can retain major gaps in spatially usable data, with more than 20% of endemic angiosperms estimated to lack a single GBIF record with valid spatial coordinates (Colli-Silva et al. 2020).

Together, these patterns highlight the difficulty of assessing data gaps among ecoregions from record numbers alone, as substantial deficiencies can persist even where occurrence data appear relatively abundant.

## 5. Conclusion and Remarkable

Overall, the results highlight that the growing availability of biodiversity occurrence data in GBIF does not necessarily translate into spatially or taxonomically representative datasets. The substantial proportion of records removed during quality filtering, the uneven representation across taxa, and the pronounced spatial concentration of records among ecoregions collectively demonstrate that the amount of data itself is an insufficient measure of data adequacy for analyses in some parts of the world.

Although the use of filters is necessary to ensure spatial consistency and reduce potential biases associated with missing or duplicated coordinates, the removal of approximately 54% of the original records inevitably reduced the amount of information available for subsequent analyses. Stricter filtering can improve the reliability and comparability of the remaining records, but it also reduces their spatial and taxonomic coverage. The observed data loss underscores the need for filtering approaches that balance quality control with the retention of informative records.

Reducing these limitations requires coordinated efforts to accelerate the digitisation and mobilisation of biological collections (James et al. 2018; Soltis et al. 2018; Eckert et al. 2024; Wilcox et al. 2026). Nevertheless, digitisation alone will not be sufficient unless biodiversity institutions also embrace standardised and interoperable infrastructures that enable broad data integration and accessibility (Wetzel et al. 2028; Sterner et al. 2020). Ultimately, sustained investments in collection digitisation, data mobilisation, and born-digital workflows are critical to transform global biodiversity databases from biased records of historical sampling efforts into more comprehensive and equitable representations of plant diversity, providing a stronger foundation for ecological research and conservation in a rapidly changing world (James et al. 2018; Pitogo et al. 2026).

## Acknowledgments

This study is part of a project (#200926/2024-1) supported by CNPq (National Council for Scientific and Technological Development), a Brazilian government agency under the Ministry of Science and Technology, through an initiative aimed at supporting early-career researchers.

